# Lineage tracing analysis of cone photoreceptor-associated cis-regulatory elements in the developing chicken retina

**DOI:** 10.1101/531475

**Authors:** Estie Schick, Sean D. McCaffery, Erin E. Keblish, Cassandra Thakurdin, Mark M. Emerson

## Abstract

During vertebrate retinal development, transient populations of retinal progenitor cells with restricted cell fate choices are formed. One of these progenitor populations expresses the Thrb gene and can be identified with the ThrbCRM1 cis-regulatory element. Short-term assays have concluded that these cells preferentially generate cone photoreceptors and horizontal cells, however developmental timing has precluded an extensive cell type characterization of their progeny. Here we describe the development and validation of a recombinase-based lineage tracing system for the chicken embryo to further characterize the lineage of these cells. The ThrbCRM1 element was found to preferentially form photoreceptors and horizontal cells, as well as a small number of retinal ganglion cells. The photoreceptor cell progeny are exclusively cone photoreceptors and not rod photoreceptors, confirming that ThrbCRM1-progenitor cells are restricted from the rod fate. In addition, specific subtypes of horizontal cells and retinal ganglion cells were overrepresented, suggesting that ThrbCRM1 progenitor cells are not only restricted for cell type, but for cell subtype as well.

## Introduction

The vertebrate retina is composed of six classes of neurons and one glial cell type that arise from multipotent retinal progenitor cells (RPCs) during development: cone photoreceptors (PRs) and rod PRs, horizontal cells (HCs), bipolar cells (BCs), amacrine cells (ACs), retinal ganglion cells (RGCs) and Müller glia. These cell types, comprising at least 100-150 subtypes^1^, are formed in overlapping temporal windows and organized into an evolutionarily conserved retinal structure^2^. The study of the molecular and genetic pathways by which cells acquire these fates is a central issue for our understanding of retinal development, and for the generation of effective therapeutic tools. In addition, due to its relative simplicity and accessibility, the retina is well-suited to serve as a microcosmic model of the developing central nervous system (CNS).

Cone PRs are critical for color and high acuity vision^3^, and their loss in diseases such as macular degeneration and retinitis pigmentosa contributes to severe visual impairment^4, 5^. Previous studies have shown through viral tracing experiments that multipotent RPCs are competent to give rise to all retinal cell types^6–8^. More recently, restricted RPCs have been identified in which daughter cells are biased to acquire specific cell fates over others, or at ratios that would not be predicted by the distribution of cell types born at that time in the retina. Some notable examples include the Ascl1 lineage, in which all cell types other than RGCs derive from RPCs marked by this bHLH factor^9^ and the Olig2 lineage. Early in development, RPCs expressing Olig2 are biased to exit the cell cycle within one or two divisions and predominantly form PRs and HCs^10^.

Similarly, there have been multiple reports of Thyroid Hormone Receptor Beta (Thrb)-related lineages in which cells preferentially choose cone PR and HC fates. An intron control region for Thrb (ThrbICR) was identified and characterized in transgenic mice, and is active in photoreceptors as well as cells in the developing inner retina^11^. In zebrafish, the Trβ2 promoter was reported to drive expression in progenitors that produced L-cone precursors and HC precursors^12^, and a Thrb cis-regulatory element (ThrbCRM1) that marks a population of likewise restricted RPCs (preferentially forming cone PRs and HCs) was reported in the chick^13^. As Thrb is the earliest known marker of developing cone PRs^14, 15^, these elements represent some of the earliest events in the cone genesis pathway. The identification of these lineages has contributed to our understanding of the gene regulatory networks that govern cone PR development, but the relationships between specific RPC populations and the particular cell types and subtypes produced are still largely unknown. Among the questions left unanswered is whether all cone PR or all HC subtypes are produced from the same RPC types.

The chicken embryo is an excellent model organism for the study of cone PR development, as cones represent the majority of PRs in the chick^16^ in contrast to a mere 3% of PRs in the mouse retina^17, 18^. However, one significant impediment in the chick system is the relative lack of established tools for the study of cell lineages. Historically, the chicken embryo has been an instrumental model for the study of development. Some of the many fundamental processes that have been pioneered and characterized in the chick include neural crest migration and development, limb development, left/right asymmetry and dorso-ventral patterning^19, 20^. However, the majority of the techniques that enabled these discoveries, such as transplantation, creation of chimeras and labelling with vital dyes, are not well-suited to the current genetics-driven nature of the field. Retroviral labelling has been utilized more recently as a method for cell lineage analyses, but its use in genetically-directed lineage tracing is limited.

An adapted method^21^ was used to describe the ThrbCRM1 population, in which a retrovirus encoding GFP carries the gp70 envelope protein, ensuring that infection is restricted to cells expressing the CAT1 receptor, which is driven by an enhancer or promoter region^13^. However, this technique does not address the remaining limitations of viral assays, namely the potential invasiveness of the viral infection, instances of spontaneous silencing^22^ the small size of marked clones, and the fact that only half of the lineage is labeled with the reporter^23^.

The advent of site-specific recombination as a mediator of genetically-directed lineage tracing has revolutionized the study of developmental biology across tissues and species^24–27^ but there is limited data about the utility and fidelity of these systems in the chicken^28, 29^. In this study, we establish a method for recombinase-based lineage tracing in the chick embryo and utilize this system to extensively characterize several cis-regulatory elements associated with the cone-related gene, Thrb. We identify discrete cell populations marked by lineage trace of these elements, suggesting the existence of multiple gene-regulatory networks involved in Thrb regulation. In addition, *in vivo* lineage tracing of the ThrbCRM1 element provides further evidence that this enhancer is active in RPCs restricted to form cone and not rod photoreceptors. Furthermore, the specific subtypes of HCs formed from this lineage are preferentially Type H1 Lim1-expressing cells, suggesting that there are genetic mechanisms restricting cell subtype as well as cell type.

## Results

### Comparison of FlpE, Cre and PhiC31 mediated recombinase-based lineage trace systems

To ensure that cells recombined using this technique will be biologically relevant, we first examined the recombination efficiency of three recombinases at basal levels in the chick. FlpE was first isolated from yeast and is widely used in Drosophila^30^, Cre which is utilized broadly in mice, and the less-commonly used PhiC31, both derive from bacteriophages^31, 32^. Retinas from embryonic day 5 (E5) chick embryos were electroporated *ex vivo* with one of three lineage trace driver plasmids, each with a recombinase placed downstream of a minimal promoter (TATA box). The retinas were co-electroporated with the respective responder plasmids in which CAG and GFP are separated by a Neomycin stop cassette flanked by FRT, LoxP, or attB/P sites (referred to as CAFNF::GFP, CALNL::GFP or CAaNa::GFP herein)^33^. Expression of the recombinase results in excision of the stop cassette from this plasmid, enabling ubiquitous expression of the reporter in those cells and in all cells deriving from them (Fig. 1a). Lastly, a broadly active UbiqC::TdTomato plasmid was included as a control for electroporation efficiency. As expected, when FlpE and PhiC31 were driven by the basal promoter, only 0.07% ± 0.02 and 0.04% ± 0.01 of electroporated cells underwent recombination, respectively (mean ± 95% CI, n=12) (Fig. 1b, d, k). However, introduction of the Cre plasmids resulted in a significant amount of recombination, with 9.4% ± 2.83 of all electroporated cells expressing GFP (mean ± 95% CI, n=12, p< 0.001) (Fig. 1c, k). This experiment was repeated and analyzed by confocal microscopy, yielding the same qualitative results (Supplementary Fig. S1).

**Figure 1.**
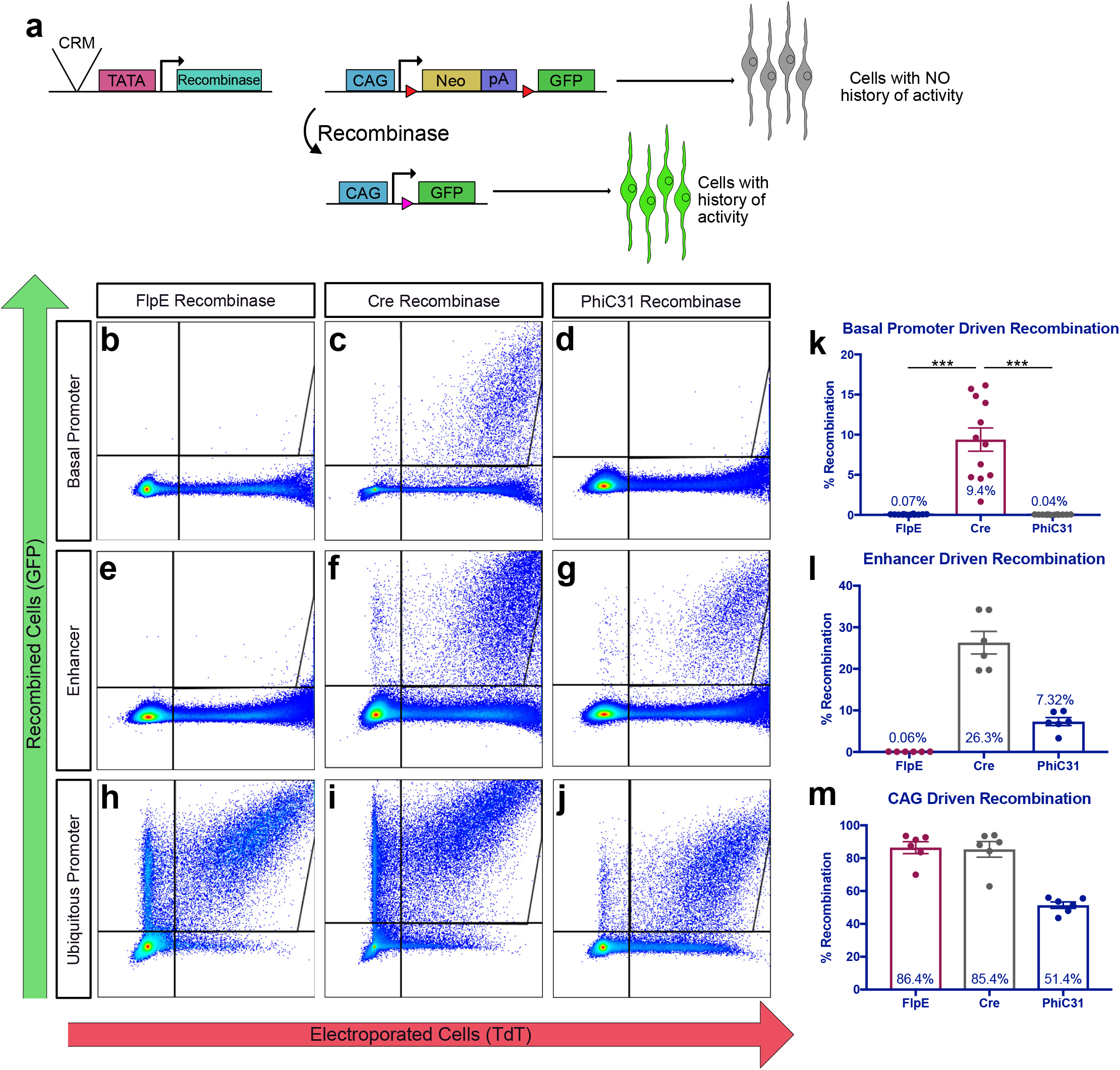
Quantification of lineage trace recombination efficiencies mediated by FlpE, Cre and PhiC31. A. Schematic representation of recombinase-based lineage tracing strategy. Red triangles represent recombinase target sequences. B-J. Representative dot plots of FACS-analyzed dissociated retinal cells electroporated with a UbiqC::TdT electroporation control, a FlpE, Cre, or PhiC31 recombinase plasmid, and the corresponding responder vector. Retinas were electroporated at E5 and harvested after two days in culture. B-D. Recombinase plasmids driven by a basal promoter. bp::FlpE (b), bp::Cre (c), bp::PhiC31 (d). E-G. Recombinase plasmids driven by the ThrbCRM1 element. ThrbCRM1::FlpE (e), ThrbCRM1::Cre (f), ThrbCR1::PhiC31 (g). H-J. Recombinase plasmids driven by the ubiquitous CAG element. CAG::FlpE (h), CAG::Cre (i), CAG::PhiC31 (j). K. Quantification of basal recombination as shown in b-d. Error bars represent 95% confidence intervals, n=12, p<0.001 upon Kruskal Wallis test with post hoc Dunn test. L. Quantification of enhancer-driven recombination as shown in e-g. Error bars represent 95% confidence intervals, n=6. M. Quantification of ubiquitous recombination as shown in h-j. Error bars represent 95% confidence intervals, n=6. CRM, cis-regulatory module; bp, basal promoter; pA, polyadenylation sequence.

Next, we determined the recombination efficiency of FlpE, Cre, and PhiC31 when driven by a cis-regulatory module (CRM). Recombinase-based lineage tracing is particularly useful for defining the lineage of cells marked by a CRM that may be active transiently during development. ThrbCRM1, an enhancer for Thrb, is active in a subset of RPCs that preferentially give rise to cone photoreceptors and HCs^13^. ThrbCRM1 was placed upstream of the TATA box in the lineage trace driver plasmids, and the resulting plasmids were electroporated into E5 chick retinas alongside the corresponding responder plasmids and the electroporation control.

Surprisingly, in retinas with ThrbCRM1::FlpE, there was no increase in recombination as compared to basal levels, with only 0.06% ± 0.01 of electroporated cells expressing GFP. In contrast, ThrbCRM1-driven Cre recombined 26.3% ± 5.34 of electroporated cells, while 7.32% ± 1.9 of cells electroporated with ThrbCRM1::PhiC31 underwent recombination (mean ± 95% CI, n=6) (Fig. 1e-g, l). Although many more cells were recombined by Cre than by PhiC31, it appears that a portion of those cells were recombined nonspecifically, as the pattern of recombined cells does not match that of ThrbCRM1 activity. The retinas electroporated with ThrbCRM1::PhiC31 show a recombination pattern consistent with ThrbCRM1 activity (Supplementary Fig. S1).

Finally, we determined maximal FlpE, Cre and PhiC31 recombination efficiencies by placing them under the control of a ubiquitous promoter. CAG was cloned upstream of the TATA box in the lineage trace driver plasmids, which were then electroporated into E5 chick retinas with the respective responder plasmids and the electroporation control. Retinas electroporated with CAG::FlpE and CAG::Cre underwent 86.4% ± 7.12 and 85.4% ± 9.28 recombination, respectively. In the case of PhiC31, 51.4% ± 3.86 of electroporated cells were recombined (mean ± 95% CI, n=6) (Fig. 1h-j, m). These results identified PhiC31 as the optimal mediator for recombinase-based lineage tracing in the chick; there is minimal basal recombination, recombination that is driven by a CRM yields a similar pattern of reporter activity to that of the CRM itself, and a majority of electroporated cells undergo recombination when PhiC31 is expressed ubiquitously.

### FlpE and Cre are suboptimal mediators of recombinase-based lineage tracing in the chick

In contrast to recombination mediated by PhiC31, and although there is minimal basal recombination with FlpE, there is no more recombination when FlpE is driven by an enhancer. Because CAG::FlpE yields recombination in more than 85% of targeted cells, we wondered whether a particular feature within CAG was contributing to this result. CAG is a hybrid construct that consists of the early CMV enhancer fused with the promoter, first exon and intron of the chicken beta-actin gene, and the splice acceptor site of the rabbit beta-globin gene^34^. As the presence of an intron has been shown to stabilize mRNA and slow its decay^35^, we thought that the intron within CAG might be stabilizing the FlpE transcript and thereby enabling FlpE-mediated recombination. This was tested by cloning an intron into the FlpE coding sequence and comparing the recombination mediated by ThrbCRM1::FlpE or ThrbCRM1:: FlpE^Intron^. The presence of the intron did not seem to have an effect, as similar amounts of GFP were detected in both conditions (Supplementary Fig. S2). To test whether the intron interfered with FlpE expression or function, the CAG element was cloned upstream of FlpE^Intron^. Robust GFP expression was observed, suggesting that the presence of the intron did not interfere with production of functional FlpE protein.

In the case of Cre-mediated recombination, the high level of basal recombination was a combined effect of leakiness from the lineage trace driver and responder plasmids. Low levels of recombination were detected in retinas when CALNL::GFP is electroporated alone, but a much higher level of recombination was observed in retinas electroporated with both plasmids (Supplementary Fig. S3). This, as well as the non-specific pattern of recombination in retinas with ThrbCRM1::Cre, indicates that of the three recombinases assessed, PhiC31 is the optimal mediator of recombinase-based lineage tracing in the developing chick embryo.

### *In vivo* Lineage Trace of Thrb CRMs

Once the PhiC31-based lineage trace system was used successfully in an *ex vivo* context, we applied the system *in vivo*. Specifically, we were interested in using this technique to examine the lineages of three CRMs for the Thrb gene. In addition to ThrbCRM1, the ThrbCRM2 and ThrbICR elements are also enhancers for Thrb^11, 13^ (Supplementary Fig. S4). Interestingly, GFP reporter expression is driven in similar patterns by the ThrbCRM1 and ThrbICR elements, with reporter-positive cells in the outer and inner retina, while ThrbCRM2 activity is restricted to the outer retina^13^. Cells marked by ThrbCRM2 activity are almost entirely also marked by ThrbCRM1 activity (Supplementary Fig. S4), and a subset of retinal cells have been reported to be marked by both ThrbCRM1 and ThrbICR activity^13^. Lineage tracing of these CRMs can provide information about the cells marked by each and may thereby facilitate a more complete understanding of their respective roles in Thrb gene regulation.

PhiC31 driver plasmids, CAaNa::GFP and CAG::nucβgal were electroporated into E3 chicken retinas in ovo. The embryos were incubated until E10, when the retina is well-organized into layers: photoreceptors are localized to the outer nuclear layer (ONL); the inner nuclear layer (INL) is arranged with HCs and amacrine cells at the apical and basal borders, respectively, and bipolar cells and Mueller glia located throughout; RGCs are positioned in the ganglion cell layer (GCL)^36, 37^. We determined the general distribution of cells by their localization and basic morphology for the three Thrb elements as well as bp::PhiC31 and CAG::PhiC31 as controls. For each condition, the quantification was averaged over a minimum of three biological replicates examined.

In retinas with the minimal promoter driving PhiC31, no electroporated cells in any layers of the retina were also GFP + (276 cells counted, n=3) (Fig. 2a, f, Supplementary Fig. S5). In retinas electroporated with CAG::PhiC31, 43.76% ± 2.65 of targeted cells in the ONL were recombined, as were 48.23% ± 9.17 and 22.93% ± 8.49 in the INL and GCL, respectively (370 cells counted, mean ± SEM, n=5) (Fig. 2b, f-g). The recombined cells were identified as PRs, HCs, ACs, RGCs and as BCs or Mueller glia, which are not easily distinguishable by localization. Additionally, there may be displaced ACs in the RGC layer that are counted as RGCs. Nonetheless, these quantifications confirm the absence of basal activity and the targeting of all cell types in the retina by electroporation.

**Figure 2.**
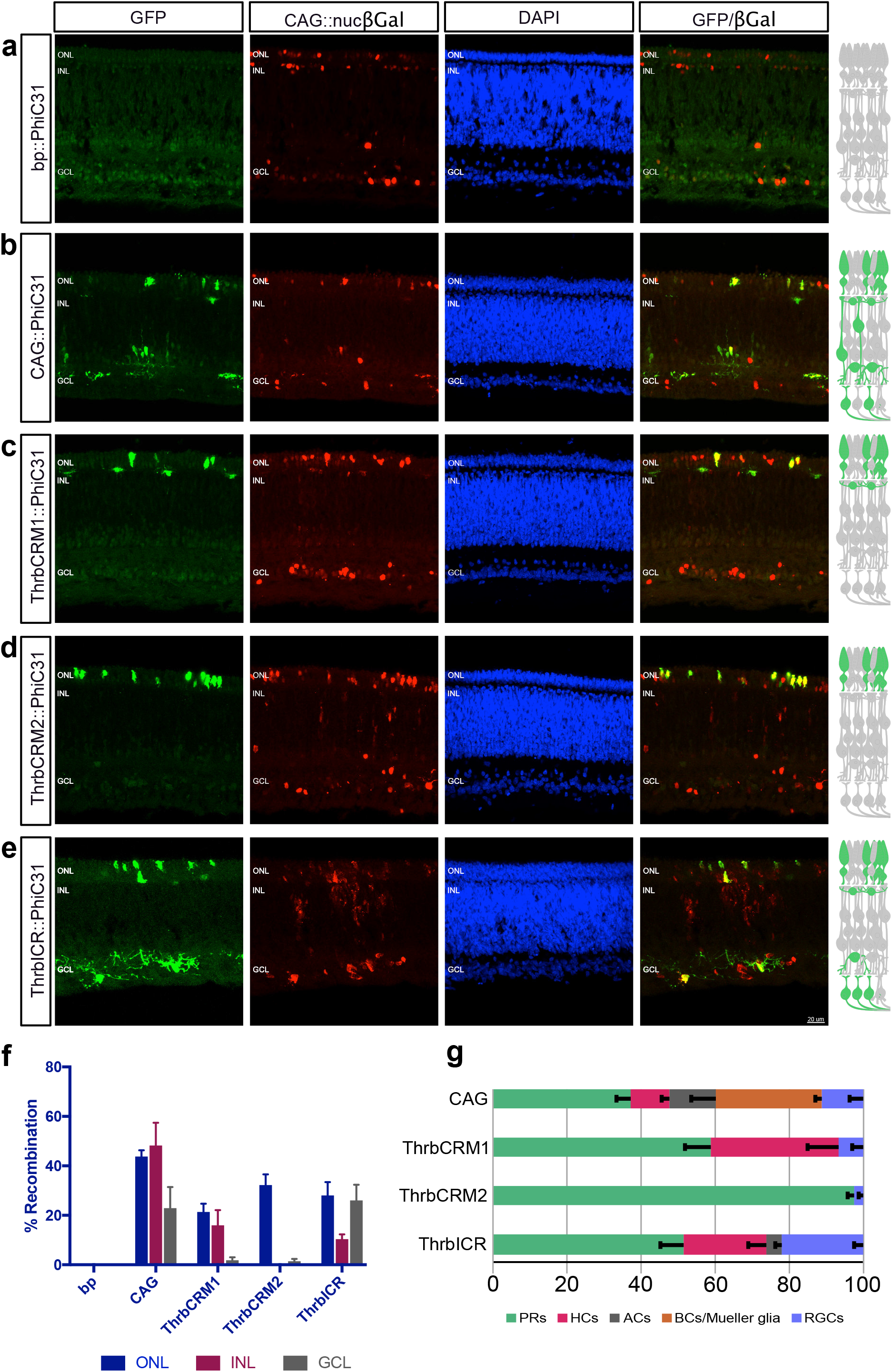
*In vivo* lineage trace of 3 Thrb CRMs yields unique patterns of activity. A-E. Representative images of vertically sectioned retinas that were electroporated *in vivo* at E3 with CAG::nucβgal as an electroporation control, bp::PhiC31 (a), CAG::PhiC31 (b), ThrbCRM1::PhiC31 (c), ThrbCRM2::PhiC31 (d), or ThrbICR::PhiC31 (e) and CAaNa::GFP. Embryos were grown until E10, and all images are maximum intensity projections. To the right of each image is a schematic of the chick retina with recombined cell types from all counted images colored in green. F. Quantification of the % recombination in each retinal layer, for each of the conditions assessed, n=3-8. G. Quantification of the % of representation for each retinal cell type among the recombined cells for that particular condition. The negative control had no recombination (f) so it was excluded here. Error bars represent SEM, n=3-8. ONL, outer nuclear layer; INL, inner nuclear layer; GCL, ganglion cell layer; bp, basal promoter; PRs, photoreceptors; HCs, horizontal cells; ACs, amacrine cells; BCs, bipolar cells; RGCs, retinal ganglion cells.

The conclusions about ThrbCRM1 activity in restricted RPCs are based on previous work in which retinas were labelled by retroviral infection such that the progeny of ThrbCRM1 + RPCs were marked with GFP, and cells with reporter expression were subsequently identified as photoreceptors or HCs by co-localization with Visinin (Rcvrn) or Lim1 (Lhx1)^13^. To confirm this lineage characterization *in vivo*, we examined the cell populations with a history of PhiC31-mediated recombination driven by ThrbCRM1 (4×40 bp core ThrbCRM1 element). 21.39% ± 3.3 of targeted cells in the ONL were recombined, as well as 16% ± 6.07 in the INL and 1.85% ± 1.16 in the GCL (1,295 cells counted, mean ± SEM, n=8). The recombined cells were primarily PRs and HCs, with a small number of RGCs (Fig. 2c, f-g).

ThrbCRM2 activity has previously been detected in a population of Visinin + cells located in the outer retina^13^. Here we traced the ThrbCRM2 lineage *in vivo*, and observed recombination in 32.25% ± 4.35 of electroporated cells in the ONL, no recombination in the INL, and 1.41% ± 0.99 recombination in the GCL (463 cells counted, mean ± SEM, n=6) (Fig. 2d, f). Almost the entirety of this population were therefore PRs, in addition to a few RGCs (Fig. 2g).

As mentioned above, the pattern of ThrbICR activity appears somewhat similar to that of ThrbCRM1, although robust ThrbICR activity was observed in RGCs in mouse^11^. Because previous work has shown that its activity can be observed as early as 6 hours post electroporation, and that a subset of cells are marked by both ThrbCRM1 and ThrbICR activity^13^, we wondered whether ThrbICR might be active in a progenitor population as well. E5 retinas electroporated with ThrbICR::GFP were pulsed with EdU for 1 hour after 1 day of culture. Multiple GFP +/ EdU + cells were detected, implying that at least a portion of ThrbICR activity is in progenitor cells (Supplementary Fig. 6). We proceeded with the *in vivo* ThrbICR lineage trace, and observed recombination in 28.04% ± 5.39 of targeted cells in the ONL, 10.35% ± 1.95 in the INL and 26.05% ± 6.31 in the GCL. These recombined cells comprise PRs, HCs, RGCs, and a small number of ACs (391 cells counted, mean ± SEM, n=5) (Fig. 2e, f-g).

### Thrb CRMs mark differing ratios of early-born retinal cells *in vivo*

The *in vivo* lineage tracing of ThrbCRM1, ThrbCRM2 and ThrbICR both supports and provides more resolution to the respective CRM activity observed *ex vivo*. In relation to the CAG::PhiC31 control, each of the three Thrb CRMs show an overrepresentation of activity in PRs, while HCs are overrepresented in the ThrbCRM1 and ThrbICR lineages but absent from the ThrbCRM2 population. RGCs marked by ThrbCRM1 and ThrbCRM2 are underrepresented relative to the CAG population, but are overrepresented in the ThrbICR lineage (Fig. 2g). To ensure that the characterization of cell types marked by each CRM is not biased by the number of cells targeted in each layer of the retina, we examined the total percentages of electroporated cells that underwent recombination in the ONL, INL, and GCL. Although the number of cells marked by electroporation varies, no trends are observed between the number of targeted cells and the percentage of recombination in each layer. (Supplementary Fig. S5).

### The ThrbCRM1 lineage includes cone photoreceptors but not rod photoreceptors

As shown above, the restricted ThrbCRM1 RPCs give rise to photoreceptors in the ONL. However, MafA (L-Maf), the earliest known marker of rods in the chick, is expressed beginning at E9, and therefore at E10 the ONL contains both cones and rods^38^. It has previously been reported that the ThrbCRM1 element is active in the mouse retina at E13.5, a period of cone genesis, but not at P0, when rods are the only photoreceptors targeted by electroporation^13^. To test if this cone vs rod specificity was evident in the chick retina, we assessed the ThrbCRM1 RPC-derived photoreceptors. However, we have observed that photoreceptors are underrepresented by CAG, which could mean that some of the photoreceptors derived from ThrbCRM1 RPCs may not be labeled with CAG-driven GFP (data not shown). To ensure that this does not impede the characterization of photoreceptors in the ThrbCRM1 lineage, we modified the PhiC31 responder vector by cloning a putative CRM for the pan-photoreceptor gene Visinin, that drives strongly in Visinin + cells, upstream of CAG (Supplementary Fig. S7). This could allow a ThrbCRM1 RPC-derived photoreceptor excluded by CAG but marked by the Visinin CRM (VisPeak) to express GFP. In an *ex vivo* context, a ThrbCRM1 lineage trace with CAaNa::GFP^VisPeak^ resulted in a slight increase of GFP + cells that were excluded from the CAG::TdT population. Additionally, more cells both within and outside of the CAG::TdT population were Visinin + when the PhiC31 responder used was CAaNa::GFP^VisPeak^ (Supplementary Fig. S7). We therefore lineage traced ThrbCRM1 at E3 using CAaNa::GFP^VisPeak^ and harvested the retinas at E10.

In order to determine whether the recombined cells in the ONL are cone photoreceptors, flat-mounted retinas were immunolabeled for Rxrg, a cone-specific marker^39^. 79.5% ± 0.74 of cells targeted by electroporation in the ONL were Rxrg + (1,834 cells counted, mean ± SEM, n=3). As Rxrg does not mark all cone photoreceptor types^40^, the subset of cells that were Rxrg - may be a cone photoreceptor subpopulation, rods, or a combination of the two. Of the targeted PRs derived from the ThrbCRM1 lineage, 92.5% ± 2.06 were Rxrg + (166 cells counted, mean ± SEM, n=3) (Fig. 3a, c). This confirms that a vast majority, if not all, photoreceptors with a history of ThrbCRM1 activity are cone photoreceptors.

**Figure 3.**
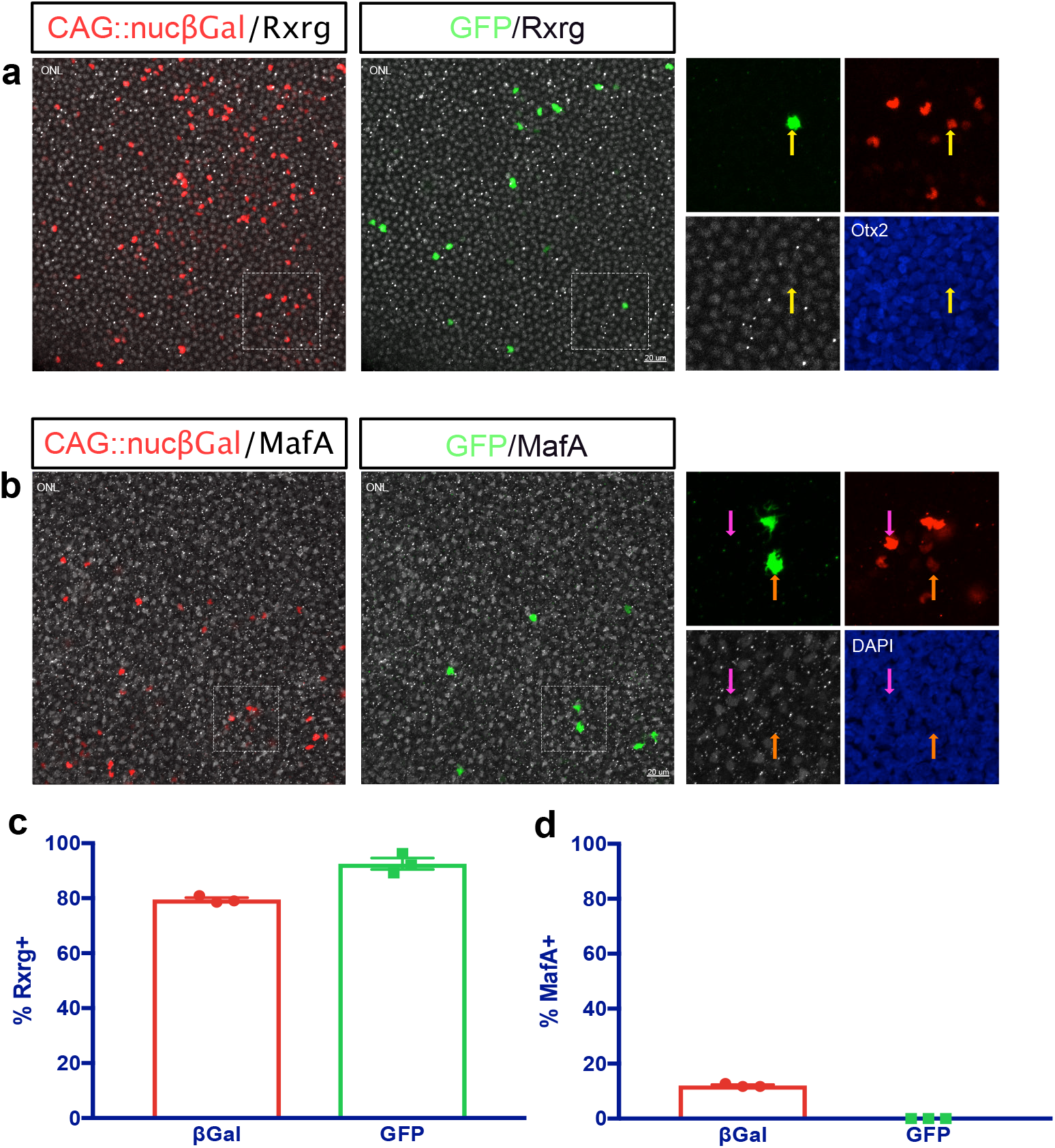
ThrbCRM1 RPCs give rise to cone PRs, but not to rods. A. Representative image of the ONL of a ThrbCRM1 lineage traced flat-mounted retina at E10, counterstained with Rxrg (white) to mark cone PRs. Areas zoomed in insets are outlined in dotted line. Yellow arrow represents electroporated GFP+/Rxrg+ cells. Electroporated βgal+ cells are shown in red, maximum intensity projection of 40x image. B. Representative image of the ONL of a ThrbCRM1 lineage traced flat-mounted retina at E10, counterstained with MafA (white) to mark rod PRs. Areas zoomed in insets are outlined in dotted line. Magenta arrow represents electroporated GFP-/MafA+ cells, orange arrow represents electroporated GFP+/MafA-cells. Electroporated βgal+ cells are shown in red, maximum intensity projection of 40x image. C. Quantification of Rxrg immunolabeled retinas as shown in a- % Rxrg+ cells of electroporated ONL cells (βgal+) and of ThrbCRM1 lineage traced ONL cells (GFP+). Error bars represent SEM, n=3. D. Quantification of MafA immunolabeled retinas as shown in b- % MafA+ cells of electroporated ONL cells (βgal+) and of ThrbCRM1 lineage traced ONL cells (GFP+). Error bars represent SEM, n=3. ONL, outer nuclear layer.

To determine whether the Rxrg - cells from the ThrbCRM1 lineage are rods or cones, we next stained retinas for MafA. 12% ± 0.35 of photoreceptors targeted by electroporation were MafA + and 88% were MafA - (1,290 cells counted, mean ± SEM, n=3). These percentages of rods and cones are similar to the previously documented numbers (14% and 86%)^16^, although the number of rods marked by MafA may be somewhat smaller given that this population is not fully formed by E10. Of the electroporated cells derived from the ThrbCRM1 RPCs, 100% were MafA - (81 cells counted, n=3) (Fig. 3b, d). This data supports the conclusion that photoreceptors in the ThrbCRM1 lineage are exclusively cone photoreceptors.

### The HCs born from ThrbCRM1 lineage are primarily the H1 type

In the chick retina, there are four types of HCs: the H1 population makes up 52% of all HCs in the retina, and is defined by expression of Lim1 and Ap2α, while the H2, H3 and H4 populations encompass the remaining HCs and are defined by Islet1 expression^41^. To characterize the HCs in the ThrbCRM1 lineage, E10 ThrbCRM1-lineage traced retinas were stained for Lim1. 58.4% (SEM±4.39) of electroporated cells in the HC layer of the INL were Lim1 + (1,510 cells counted, mean ± SEM, n=3). This is roughly equivalent to the percentage of H1 HCs in the chick retina. Of the cells derived from the ThrbCRM1 lineage, an overwhelming 88.6% ± 3.52 were Lim1 + (124 cells counted, mean ± SEM, n=3) (Fig. 4a, c).

**Figure 4.**
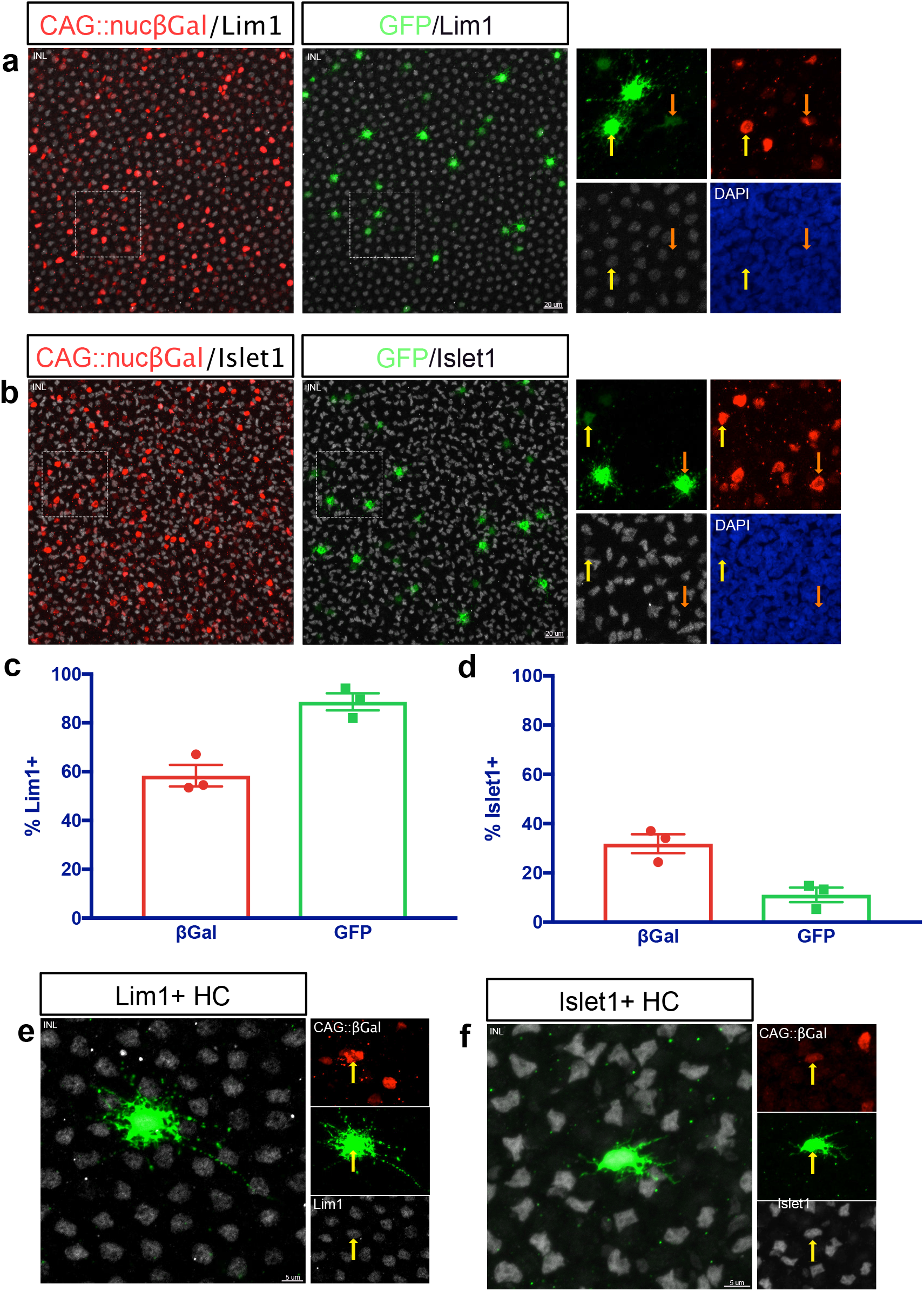
The HCs in the ThrbCRM1 lineage are predominantly the H1 type. A. Representative image of the INL of a ThrbCRM1 lineage traced flat-mounted retina at E10, counterstained with Lim1 to mark H1 HCs. Areas zoomed in insets are outlined in dotted line. Yellow arrow represents electroporated GFP+/Lim1+ cells, orange arrow represents electroporated GFP+/Lim1-cells. Maximum intensity projection of 40x image. B. Representative image of the INL of a ThrbCRM1 lineage traced flat-mounted retina at E10, counterstained with Islet1 to mark H2, H3 and H4 HCs. Areas zoomed in insets are outlined in dotted line. Yellow arrow represents electroporated GFP+/Islet1+ cells, orange arrow represents electroporated GFP+/Islet1-cells. Maximum intensity projection of 40x image. C. Quantification of Lim1 immunostained retinas as shown in a- % Lim1+ cells of electroporated cells in the HC layer of the INL (βgal+) and of ThrbCRM1 lineage traced INL cells (GFP+). Error bars represent SEM, n=3. D. Quantification of Islet1 immunostained retinas as shown in b- % Islet1+ cells of electroporated cells in the HC layer of the INL (βgal+) and of ThrbCRM1 lineage traced INL cells (GFP+). Error bars represent SEM, n=3. E-F. Representative images of Lim1+ (e) or Islet1+ (f) HCs within the ThrbCRM1 lineage. INL, inner nuclear layer; HC, horizontal cell.

The approximately 12% of Lim1 - HC cells are predicted to represent the H2, H3 and H4 cell types. To confirm this, additional E10 ThrbCRM1-lineage traced retinas were immunostained for Islet 1, which marks these HC populations. As expected, 11.1% of the HCs within the ThrbCRM1 lineage were Islet1 +, while 88.9% ± 2.96 were Islet1 - (202 cells counted, mean ± SEM, n=3) (Fig. 4b, d). This confirms that in addition to the bias of ThrbCRM1 progenitors to form HCs over other cell types, those HCs are primarily of the H1 subtype.

The morphologies of the different HC classes have been characterized extensively in the chick, with the H1 type defined as having an axon and a narrow field of thick and short dendrites with bulbous endings. The H2-H4 types have no axon and have thin, irregular dendrites^41, 42^. The Lim1 + (H1) and Islet1 + (H2, H3 or H4) HCs in the ThrbCRM1 lineage match those descriptions (Fig. 4e-f).

### A small population of RGCs is derived from ThrbCRM1 progenitors

As shown above, the ThrbCRM1 lineage also includes a small number of RGCs (Fig. 2g). We determined that 0.91% ± 0.28 of electroporated RGCs in flat-mounted retinas were part of the ThrbCRM1 lineage (1,376 cells counted, mean ± SEM, n=3) (Fig. 5a, d). Although these RGCs account for a small subset of the ThrbCRM1 lineage, we were interested in characterizing these cells further. E10 retinas lineage traced for ThrbCRM1 were immunolabeled for Brn3a (Pou4f1), which marks a class of RGCs^43^, and 35.29% ± 6.19 were Brn3a+(46 cells counted, mean ± SEM, n=3) (Fig. 5c). Considering that in the mouse retina, around 70% of GCs are Brn3a +, half of which co-express Brn3b^43^, Brn3a may be underrepresented in the ThrbCRM1 lineage. In fact, Brn3a + RGCs make up 68.5% ± 5.48 of electroporated cells in the GCL (n=1,192 cells counted, mean ± SEM, n=3). The Brn3a - cells are assumed to be Brn3b + and/or Brn3c +, but the lack of functional antibodies for these chick proteins did not allow for this confirmation.

**Figure 5.**
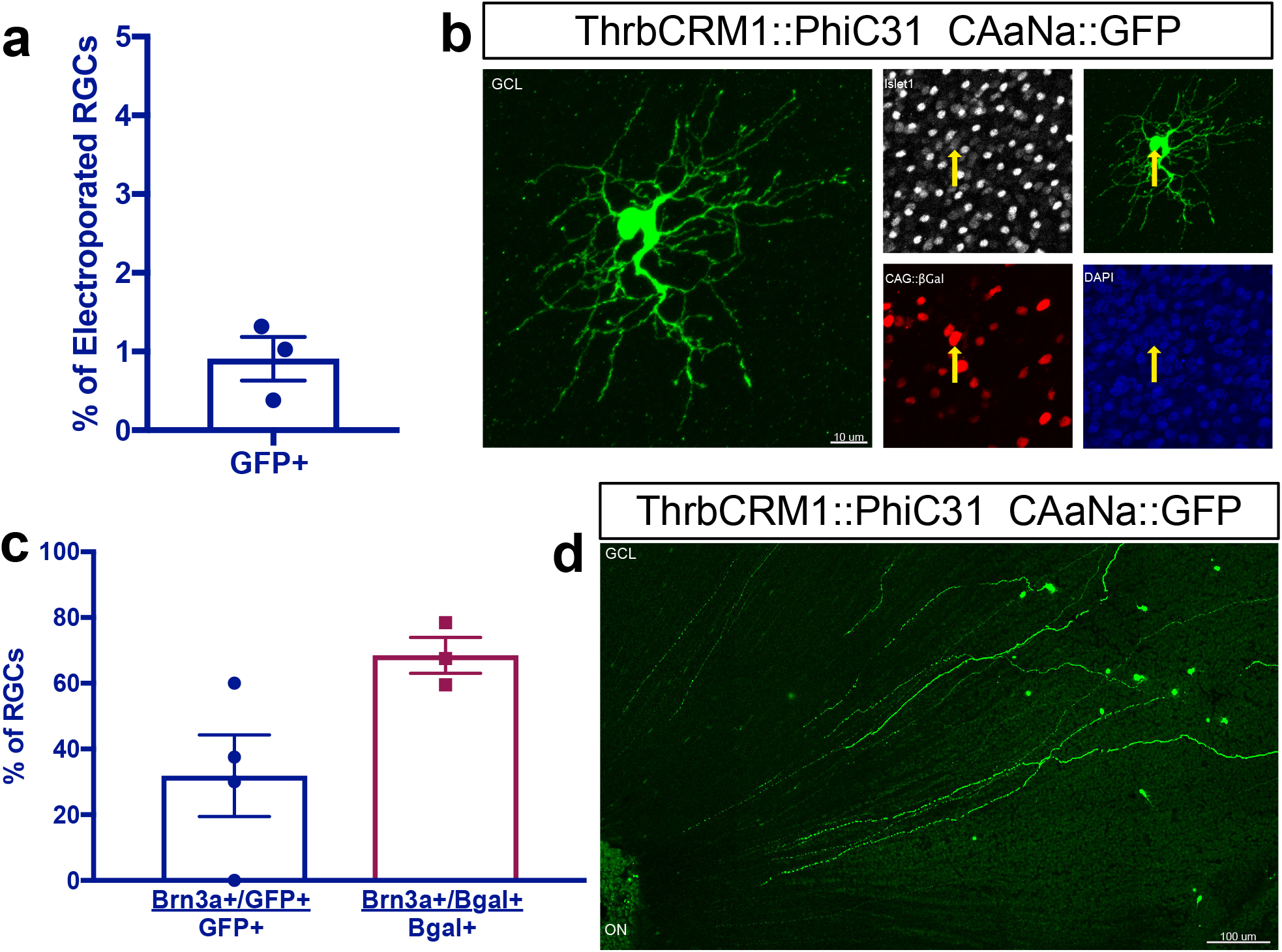
A small number of RGCs are produced from ThrbCRM1 RPCs, and the majority are not Brn3a+. A. Quantification of the percentage of electroporated cells in the GCL that were recombined in E10 ThrbCRM1 lineage traced flat-mounted retinas. Error bars represent SEM, n=3. B. Representative image of Islet1+ RGC from the ThrbCRM1 lineage. Yellow arrows show an electroporated GFP+/Islet1+ RGC. Maximum intensity projection of 40x image. C. Quantification of the % of ThrbCRM1 RGCs that are Brn3a+, and of the % of electroporated RGCs that are Brn3a+ in E10 ThrbCRM1 lineage traced flat-mounted retinas. Error bars represent SEM, n=3-4. D. Tiled image of RGCs derived from ThrbCRM1 RPCs, with axons reaching the optic nerve head, labeled as ON. Maximum intensity projection of tiled image. RGCs, retinal ganglion cells; GCL, ganglion cell layer; ON, optic nerve.

In addition, although there has not been a thorough molecular characterization of chick RGC subtypes, previous studies have classified RGCs in the chick embryo into 4 groups by general morphology^44^. Group 1 RGCs have small soma with a narrow, localized dendritic field; Group 2 RGCs have mid-sized cell bodies with medium-sized dendritic fields; Group 3 RGCs have medium-sized cell bodies and wide dendritic fields; and Group 4 RGCs have large soma and a wide, extensively branched, dendritic field. Although development is still ongoing at E10, we can identify cells among the ThrbCRM1 RPC-derived RGCs that may correspond to Group 3 RGCs (Fig. 5b).

## Discussion

*In vivo* recombinase-based lineage tracing is a powerful method for determining the cell types with a history of cis-regulatory activity during development. To be effective, a system with a low level of basal promoter activity and a high efficiency of recombination is optimal. Low basal promoter activity allows for reporter-positive cells to be ascribed to the activity of a cis-regulatory element, and is an essential parameter that needs to be empirically determined for a given paradigm. The Stagia3-based heterologous promoter used here has been shown previously to have low basal activity in GFP reporter assays in the chick retina^45, 46^ and in this study, has low basal activity with the use of PhiC31 and FlpE, but not with Cre. However, a previous study successfully used the same Cre plasmid in the developing mouse retina, suggesting that the cellular context is important^33^. In addition, it is likely that other configurations of Cre plasmids could be generated to tighten its expression and make it effective in contexts where this Cre plasmid is not. An additional factor that affects basal activity is influence of other genetic components. A benefit of this plasmid-based system is that the reporter and recombinase DNA remain episomal and are not subject to integration effects such as epigenetic silencing or aberrant activation, which components such as the Tol2 system could be subject to. The downside is that plasmids are diluted with divisions and the signal therefore becomes weaker in marked cells. A second element required for effective lineage systems is high recombination efficiency. In comparisons of CAG-driven recombination efficiency, the PhiC31 system was less effective compared to the Cre and FlpE systems. Thus, it is likely that the PhiC31 *in vivo* lineage tracing system is underrepresenting, to some extent, the percentages of cells labeled by a given cis-regulatory element. Further modification of the PhiC31 system or those based on Cre and FlpE to make them more effective would be useful for simultaneous dual-recombinase experiments.

The Thrb gene has been a target of multiple cis-regulatory studies due to its role in early cone genesis^11–13^. Interestingly, all three of these studies identified activity of Thrb elements in cells other than cone photoreceptors. In chicken, the ThrbCRM1 element was shown to be expressed in RPCs that primarily generate cone photoreceptors but also HCs^13, 47^. A larger genomic element in zebrafish was also found to be active in cones, but live imaging experiments identified reporter-positive RPCs that formed cones as well as HCs and RGCs from a prior division^12^. In mice, the ThrbICR element was used in a transgenesis assay and was found to label cones as well as HCs and RGCs^11, 13^. These results can be explained either by activity of these elements initiated in restricted RPCs that generate combinations of these cell types or by activation of these elements in postimitotic cells. The use of live imaging and targeted retroviral labeling in zebrafish and chicken, respectively, both suggest that the restricted RPC model likely accounts for much of this linked expression. We postulate the following model for the relationship of cis-regulatory activity with cell types, using brackets to define the cell types predominantly formed from an RPC type (Fig. 6). Multipotent RPCs divide asymmetrically to generate another multipotent RPC and a RPC[G,C,H] with activity of the ThrbICR element. Division of the RPC[G,C,H] generates a postmitotic RGC (currently, a dedicated RGC RPC state has not been identified) and an RPC[C,H] defined by the ThrbCRM1 element. Divisions of this RPC primarily generate cones and horizontal cells which in fish, at least, are likely to have dedicated or homotypic RPCs (RPC[C] and RPC[H]^48^) that generate these cells. Lastly, the ThrbCRM2 element is almost exclusively active in a subset of photoreceptors that are presumed to be cones as they are formed prior to the start of rod photoreceptor genesis and they are specifically segregated from photoreceptors with active Rhodopsin promoter^46^. This model is consistent with the live imaging observations of clone divisions observed in zebrafish^12^ and the cumulative cell type distributions observed here.

**Figure 6.**
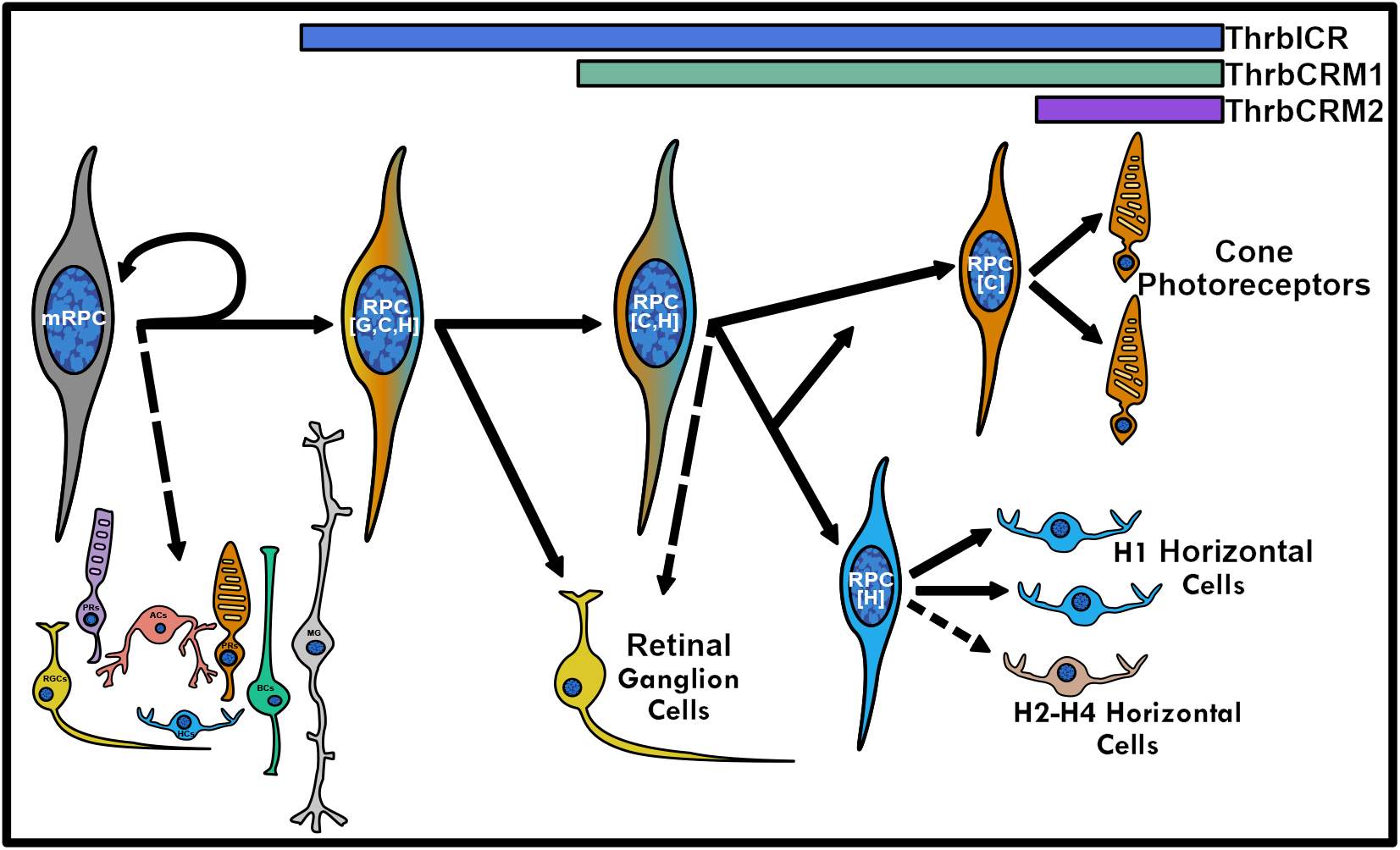
A proposed model for the relationship between Thrb cis-regulatory activity and the generation of early-born retinal cell types.

Additional clonal analyses performed in zebrafish have shown that RGCs are frequently born from divisions in which an RPC forms one proliferating cell and one differentiating cell, and are rarely born alongside another RGC^49^. However, the clone composition observed was far more diverse than the lineages reported by Suzuki et al, and of those described here, both of which utilized genetically-directed lineage analyses to specifically examine transient progenitor populations. It is therefore likely that these studies accessed a restricted lineage, but that many other RPC divisions are not restricted. In addition, we don’t know the extent to which ThrbCRM1 RPCs, or other restricted RPCs associated with Thrb, are stochastic within a set of defined choices.

While the analysis of the mouse ThrbICR transgenic did not show evidence of activity in RPCs using BrdU labeling, these experiments were conducted at E14.5, which could be past the developmental time window of an RPC[G,C,H]. Additionally, the same study did not identify βgal reporter expression in HC cells, however, HC markers were not explicitly examined and HCs are localized to the inner retina with RGCs during embryonic development. Our model of the ICR activity in RPC[G,C,H]s is based on the zebrafish data and the ThrbICR activity in RPCs shown here, but we can not yet exclude that the RGC activity is independently activated in postmitotic RGCs.

The HC population lineage-traced *in vivo* by ThrbCRM1 was composed of almost 90% Lim1 + Type I HCs, in contrast to the 52% that would be predicted from the total population frequencies. This strongly suggests a model in which ThrbCRM1 RPCs are restricted not only with regard to the types of cells that they generate (cones and HCs, primarily), but also the specific cell subtype (Type I HCs). In addition, there was a small population of RGCs that lineage-traced from the ThrbCRM1 population, and that were not previously identified in analysis of the ThrbCRM1 element. This could be due to the very low number of RGCs, but also to the timing of the studies, as the current lineage tracing study introduced DNA two days prior to the E5 electroporation used previously. While several studies have identified genetically encoded reporters that have been linked to restricted RPC expression, only two have implicated the formation of a specific cell subtype from heterotypic RPCs^12, 50^. One of these is the previously mentioned study in zebrafish in which a large cis-regulatory element with regions 5’ and 3’ to the first exon of Trβ2 gene drove reporter expression solely in L-cones. However, a recombinase strategy was not utilized and so it is unclear if this reporter expression represents transcriptional activity of the reporter from an L-cone specific element or from reporter expression inititated in the RPCs, as was concluded. The zebrafish element encompassed 9kb of regulatory DNA and a region homologous to the photoreceptor-specific chicken ThrbCRM2 element was present in the zebrafish construct. Thus, it is possible that this L-cone expression was initiated independently in postmitotic cones (or in homotypic cone RPCs) by this cis-regulatory region. It will be interesting to examine the cone subtypes with a history of ThrbCRM2 and ThrbCRM1 activity.

## Methods

### Animals

All experimental procedures involving animals were carried out in accordance and in consultation with the City College of New York Institutional Animal Care and Use Committee for the use of early stage embryonated avian eggs. Fertilized chick eggs were obtained from Charles River, stored at 16 °C for 0-10 days and incubated in a 38 °C humidified incubator.

### DNA Plasmids

CALNL::GFP, CAFNF::GFP, pCAG::FlpE, pCAG::Cre and pCAG::GFP^33^, UbiqC::TdT^51^, bp::Cre^52^, ThrbCRM1(4X)::GFP and ThrbICR::GFP^13^, and CAG::IRFP^53^ were described previously, and CAG::nucβgal (Cepko Lab, Harvard Medical School) was obtained from the Cepko lab. CAG::TdT was made by ligating a CAG fragment excised from pCAG::GFP with Sal1/EcoR1 into a Sal1/EcoR1 digested Statia plasmid^46^. To generate bp::FlpE, EGFP was removed from Stagia3^45^ with Age1/BsrG1 and blunt ends were generated with DNA Polymerase I, Large (Klenow) Fragment (NEB, M0210S). The FlpE fragment was excised from pCAG::FlpE using Not1/EcoR1 and blunt ends were created in the same way. The blunt-ended FlpE fragment was then ligated into the blunt-ended Stagia3 plasmid. Bp::PhiC31 was made by PCR-amplifying the PhiC31 coding region from pPhiC31o^54^ (Addgene plasmid #13794) using an Xma1-tagged forward primer and a BsrG1-tagged reverse primer (see Supplementary Table S1). The digested PCR product was ligated into an Age1/BsrG1 digested Stagia3 plasmid. A Sal1/EcoR1 digested CAG fragment from pCAG::GFP was cloned into a Sal1/EcoR1 digested bp::PhiC31 plasmid to generate CAG::PhiC31. The ThrbCRM1::FlpE, ThrbCRM1::Cre and ThrbCRM1::PhiC31 plasmids were created by ligation of a ThrbCRM1 element flanked by Sal1 and Xho1 sites^13^ into a bp::FlpE plasmid digested with Sal1, a bp::Cre plasmid digested with Sal1/Xho1 or a bp::PhiC31 plasmid digested with Sal1. ThrbCRM1(4X)::PhiC31 was made by cloning a Not1/EcoR1 digested 4X ThrbCRM1 fragment from ThrbCRM1(4X)::GFP into a Not1/EcoR1 digested Bp::PhiC31 plasmid. ThrbCRM2::PhiC31 and ThrbICR::PhiC31 were made by inserting the relevant Thrb regulatory elements^13^ into Bp::PhiC31 with Sal1/Xho1 or EcoR1, respectively.

To clone the CaANa::GFP plasmid, primers (attPLongNeoF1 and attBLongNeoR1) containing minimal attB and attP site sequences^55^ were used to PCR-amplify the Neomycin stop cassette from the CALNL::GFP plasmid. This PCR product contained the Neomycin stop cassette flanked by partial attB and attP sites, and was cloned into pGemTeasy (Promega, A1360). The pGem plasmid was used as a template for a subsequent PCR with attPLongNeoF2 and attBLongNeoR2 primers (see Supplementary Table S1), generating a PCR product in which the Neomycin stop cassette is flanked by complete attP and attB sites which are in turn flanked by Xho1 sites. This fragment was digested with Xho1 and cloned into the XhoI digested CALNL::GFP vector, replacing the loxP-flanked stop cassette.

To make the ThrbCRM1::FlpE^Intron^ plasmid, a Geneblock (IDT) was designed with a partial FlpE sequence followed by a 90 bp Beta-actin intron sequence surrounded by conserved CAGG sites, and another partial FlpE sequence (see Supplementary Table S2). The Geneblock was PCR-amplified and sequentially digested with AgeI and SwaI before ligation into an Age1/Swa1 digested ThrbCRM1::FlpE plasmid. CAG::FlpE^Intron^ was then made by cloning a Sal1/EcoR1 digested CAG fragment from CAG::FlpE into the correspondingly digested ThrbCRM1::FlpE^Intron^ plasmid, replacing the ThrbCRM1 element with CAG. To make VisPeak::GFP, the putative Visinin CRM (chrUn_NT_470806v1:2,267-2,828) was identified upon analysis of ATAC-Seq data (Sruti Patoori and Mark Emerson, unpublished observations) aligned to the chicken genome (Galgal5) in UCSC Genome Browser, and PCR-amplified from chicken genomic DNA (see Supplementary Table S1). The amplicon was cloned into PGemTeasy and sub-cloned into Stagia3 with EcoR1. To make VisPeak::PhiC31, the VisPeak element was cloned from VisPeak::GFP into Bp::PhiC31 with EcoR1 and to make CAaNa::GFP^VisPeak^, it was PCR amplified from VisPeak::GFP with primers containing Bcu1 sites (see Supplementary Table S1), digested with Bcu1 and cloned upstream of CAG into a Bcu1 digested CAaNa::GFP plasmid. All of the plasmids described above were verified by restriction digestion and sequencing.

### DNA Electroporation and Culture

All *ex vivo* and *in vivo* electroporations were carried out as described previously^52^ using a Nepagene NEPA21 Type II Super Electroporator. DNA mixes for *ex vivo* electroporations contained 100 ng/μl of all plasmids with UbiqC or CAG promoters and/or FlpE, Cre and PhiC31, and 160 ng/μl of all other plasmids. Retinas were dissected at E5 and cultured for 1-2 days upon electroporation, as described previously^13^. For *in vivo* electroporations, the DNA mixes included 1.5 μg/μl of all plasmids. Retinas in E3 embryos were electroporated and incubated in ovo until E10, at which point the electroporated patches of the retina were dissected.

### Retina Dissociation and Flow Cytometry

Any remaining retinal pigment epithelium and the condensed vitreal matter were dissected from cultured retinas in HBSS (GIBCO, 14170112) and the retinas were then dissociated with a papain-based protocol as described previously (Worthington, L5003126)^56^. Retinal cells used to determine recombinase efficiency were subsequently fixed in 4% paraformaldehyde for 15 minutes, washed 3X in 1XPBS and filtered into 4 mL FACS tubes (BD Falcon, 352054) through 40 μm cell strainers (Biologix, 15-1040). Cells were analyzed with a BD LSR II flow cytometer using 488 and 561 nm lasers. All experiments included control retinas that were either non-electroporated, electroporated with CAG::GFP or with UbiqC::TdT, and were used to generate compensation controls and to define gates. All FACS data was analyzed with FlowJo Version 10.2.

### Immunohistochemistry and EdU labelling

Cultured retinas were fixed in 4% paraformaldehyde for 30 minutes, washed 3X in 1XPBS, and cryoprotected with 30% sucrose/0.5XPBS. Retinas were flash-frozen in OCT (Sakura Tissue-Tek, 4583), and 20 μm vertical sections were acquired with a Leica cryostat and collected on slides (FisherBrand, 12-550-15). All immunofluorescence staining of slides was performed as described previously^52^.

The primary antibodies used were: chicken anti-GFP (ab13970, Abcam, 1:2000), rabbit anti-GFP (A-6455, Invitrogen, 1:2000), chicken anti-β-galactosidase (ab9361, Abcam, 1:1000), mouse IgG1 anti-β-galactosidase (40-1a-s, DSHB 1:20), mouse IgG1 anti-Visinin (7G4-s, DSHB, 1: 500), rabbit anti-MafA (gift from Celio Pouponnot, 1:400)^57^, mouse IgG2b anti-Rxrg (SC-365252, Santa Cruz Biotechnology, 1:100), mouse IgG1 anti-Lim1 (4F2-c, DSHB, 1:30), mouse IgG1 anti-Islet1 (39.3F7, DSHB, 1:100), mouse IgG2b anti-Islet1+2 (39.4D5, DSHB, 1:10), mouse IgG1 anti-Brn3a (MAB1585, EMD Millipore, 1:800), and rabbit anti-Otx2 (AB21990, Abcam, 1:500). All secondary antibodies were obtained from Jackson Immunoresearch and were appropriate for multiple labeling. Alexa 488- and 647-conjugated secondary antibodies were used at 1:400 and cy3-conjugated antibodies were used at 1:250. A 1 μg/μl 4’6-diamidino-2-phenylindole (DAPI) solution was applied to the slides for nuclear staining prior to mounting in Fluoromount-G (Southern Biotech, 0100-01) with 34×60 mm coverslips (VWR, 48393-106).

Flat-mount staining of dissected patches from retinas electroporated *in vivo* was carried out as described previously^58^, with some modifications. Dissected patches from retinas of the same condition were placed in a single well of a 24-well plate (Corning, CLS3524) and blocked in sterile-filtered 1XPBS with 0.5% Tween (VWR, 97062-332) and 10% serum overnight at 4 °C. Primary antibodies were added at the concentrations listed above in sterile-filtered 1XPBS with 0.5% Tween and 10% serum, and retinas were placed on a shaker at room temperature for 1 hour and then placed at 4 °C for 3 nights. Retinas were washed on a shaker 3X30 minutes with sterile filtered 1XPBS with 0.5% Tween. Secondary antibodies were added at the concentrations listed above in sterile-filtered 1XPBS with 0.5% Tween, and retinas were placed at 4 °C for 2 nights. Retinas were washed on a shaker 3X 30 minutes with sterile-filtered 1XPBS and stained with a 1 μg/μl DAPI solution on a shaker for 1 hour at room temperature. Retinas were mounted as described above on slides that had been bordered with a liquid blocker.

For EdU labeling, retinas were incubated in culture media with 50 μM Edu for 1 hour and fixed as described above. EdU detection was performed with a Click-iT EdU Alexa Fluor 647 imaging kit (Invitrogen, C10340).

### Imaging and Image Processing

All confocal images of vertically sectioned or flat-mount retinas were acquired with a Zeiss LSM710 inverted confocal microscope and ZEN Black 2015 21 SP2 software. Images were acquired at 1024 × 1024 resolution with an EC Plan-Neofluar 40x/1.30 Oil DIC M27 objective or an EC Plan-Neofluar 63x/1.25 Oil MIC objective. Images were analyzed with FIJI^59^. For retinas imaged in vertical section, Z-stacks from a minimum of three (maximum of 9) retinas in each condition were counted using the Cell Counter plugin for ImageJ. For whole-mounted retinas, three Z-stacks each from three retinas per condition were counted. All counts from the three technical replicates were averaged to a single data point and used for mean and SEM calculations. Images of whole retinas were acquired with an AxioZoom V16 microscope using a PlanNeoFluar Z 1x objective. All figures were assembled using the Affinity Designer vector editing program, and any adjustments to brightness and contrast were applied uniformly.

### Statistical Analysis

Graphs were made using Graphpad Prism7 software or Excel, and error bars represent 95% Confidence Interval or SEM, as noted. A Shapiro-Wilk Normality test and a Levene’s Homogeneity of Variances test were performed in RStudio version 1.1.447 and the results were significant. Therefore, a non-parametric Kruskal Wallis test with a post hoc Dunn test (adjusted with the Benjamin-Hochberg method) was calculated using RStudio.

### Data Availability

No datasets were generated or analyzed during the current study.

## Acknowledgements

Support was provided by National Science Foundation grant CAREER 1453044 (to MME) and NIMHD 3G12MD007603-30S2 (CCNY). The content is solely the responsibility of the authors and does not necessarily represent the official views of the National Institute On Minority Health and Health Disparities or the National Institutes of Health or the National Science Foundation. Jeffrey Walker, Andre Washington, and Jorge Morales provided excellent technical support with flow cytometry, cryosectioning, and confocal microcopy experiments. Celio Pouponnot kindly provided the MafA antibody. The 7G4, 40-1a, 4F2, 39.3F7, and 39.4D5 antibodies developed by Suzanne Bruhn/Constance Cepko, Joshua Sanes, and Susan Brenner-Morton/Thomas Jessell were obtained from the Developmental Studies Hybridoma Bank developed under the auspices of the NICHD and maintained by the Department of Biology at the University of Iowa. We thank Miruna Ghinia for advice on flat-mount imaging of retinas, Sruti Patoori for providing access to unpublished ATAC-Seq data and the members of the Emerson lab for support throughout the project.

## Author Contributions

ES, SDM and MME conceived the study and designed experiments. ES, SDM, EEK, CT and MME performed the experiments. ES and MME analyzed the data and prepared the manuscript. The manuscript was reviewed and edited by all authors.

