## Supplements for "Lineage tracing analysis of cone photoreceptor-associated cis-regulatory elements in the developing chicken retina"

**Supplementary Figure S1.** Qualitative assessment of lineage trace recombination efficiencies mediated by FlpE, Cre and PhiC31.
A-I. E5 retinas were electroporated with CAG::nucβgal, the recombinase plasmid shown on the left, and the appropriate responder plasmid. Retinas were harvested after 2 days in culture and imaged by confocal microscopy for βgal (red), GFP (green) and DAPI (blue).
A-C. Representative images of basal recombination in retinas electroporated with bp::FlpE (a), bp::Cre (b) or bp::PhiC31 (c).
D-F. Representative images of enhancer-driven recombination in retinas electroporated with ThrbCRM1::FlpE (d), ThrbCRM1::Cre (e) or ThrbCRM1::PhiC31 (f).
G-I. Representative images of ubiquitous recombination in retinas electroporated with CAG::FlpE (g), CAG::Cre (h) or CAG::PhiC31 (i).
J. Representative image of ThrbCRM1 enhancer activity. Retinas were electroporated at E5 with ThrbCRM1::GFP and CAG:: nucβgal, and harvested after two days in culture. Maximum intensity projection of 40x image.
OR, outer retina; IR, inner retina; bp, basal promoter; Enh, enhancer.

**Supplementary Figure S2.** Effects of insertion of an intron into FlpE on FlpE activity.
A-B. Representative images of whole retinas electroporated with CAG::mCherry as an electroporation control, ThrbCRM1::FlpE (a) or ThrbCRM1::FlpE$^{Intron}$ (b), and CAFNF::GFP at E5 and fixed after two days in culture.
C-D. Representative images of whole retinas electroporated with CAG::mCherry as an electroporation control, CAG::FlpE (c) or CAG::FlpE$^{Intron}$ (d), and CAFNF::GFP at E5 and fixed after two days in culture.

**Supplementary Figure S3.** Contribution of leaky bp::Cre and CALNL::GFP to basal recombination levels.
A-C. Representative images of whole retinas electroporated with CAG::mCherry as an electroporation control, bp::Cre (a), CALNL::GFP (b), or bp::CRE in combination with CALNL::GFP (c) at E5 and fixed after two days in culture.

**Supplementary Figure S4.** Alignment of Thrb enhancers to the chick genome.
A. Alignment of ThrbCRM1, ThrbCRM2 and ThrbICR (originally described in mouse) elements to the Galgal5 genome in UCSC Genome Browser. The Trβ2 isoform is shown in full.
B. Quantification of the % of ThrbCRM2 cells that are in the ThrbCRM1 population from FACS analyzed retinal cells. Error bars represent SEM, n=3.
OR, outer retina; IR, inner retina.

**Supplementary Figure S5.** Quantification of *in vivo* lineage tracing of Thrb regulatory elements.
A. Quantification of total number of electroporated cells counted in each of the conditions assessed (as shown in Figure 2a-e). N=3-8.
B. Quantification of the total number of electroporated cells counted per retinal layer in each of the conditions assessed. N=3-8.
C. Quantification of overall % recombination in each condition assessed (Total GFP/Total βgal + GFP only). Error bars represent SEM, n=3-8.

ONL, outer nuclear layer; INL, inner nuclear layer; GCL, ganglion cell layer; bp, basal promoter.

**Supplementary Figure S6.** The ThrbICR element is active in some dividing cells.
A. Representative confocal image of a retina electroporated with CAG::nucβgal as an electroporation control, and ThrbICR::GFP at E5, pulsed with EdU for 1 hour after 1 day in culture, and immediately fixed. Images are maximum intensity projections, zoomed insets are single z-planes. Yellow arrow represents electroporated GFP+/EdU+ cells.
OR, outer retina; IR, inner retina.

**Supplementary Figure S7.** Quantitative assessment of CAaNa::GFP modified with VisPeak.
A. Representative image of a retina electroporated with VisPeak::GFP plasmid at E5, fixed after 2 days in culture, and counterstained with Visinin. Merge shows extensive colocalization of Visinin with GFP+ cells.
B. Quantification of FACS analyzed retinal cells, electroporated at E5 with CAG::TdT, ThrbCRM1::PhiC31, and CAaNa::GFP or CAaNa::$GFP^{VisPeak}$ and dissociated and fixed after two days in culture. Error bars represent SEM, n=3.
C. Quantification of basal recombination in E10 retinas electroporated *in vivo* at E3 with bp::PhiC31 and CAaNa::$GFP^{VisPeak}$.
OR, outer retina; IR, inner retina; bp, basal promoter.

### Supplementary Figure S1

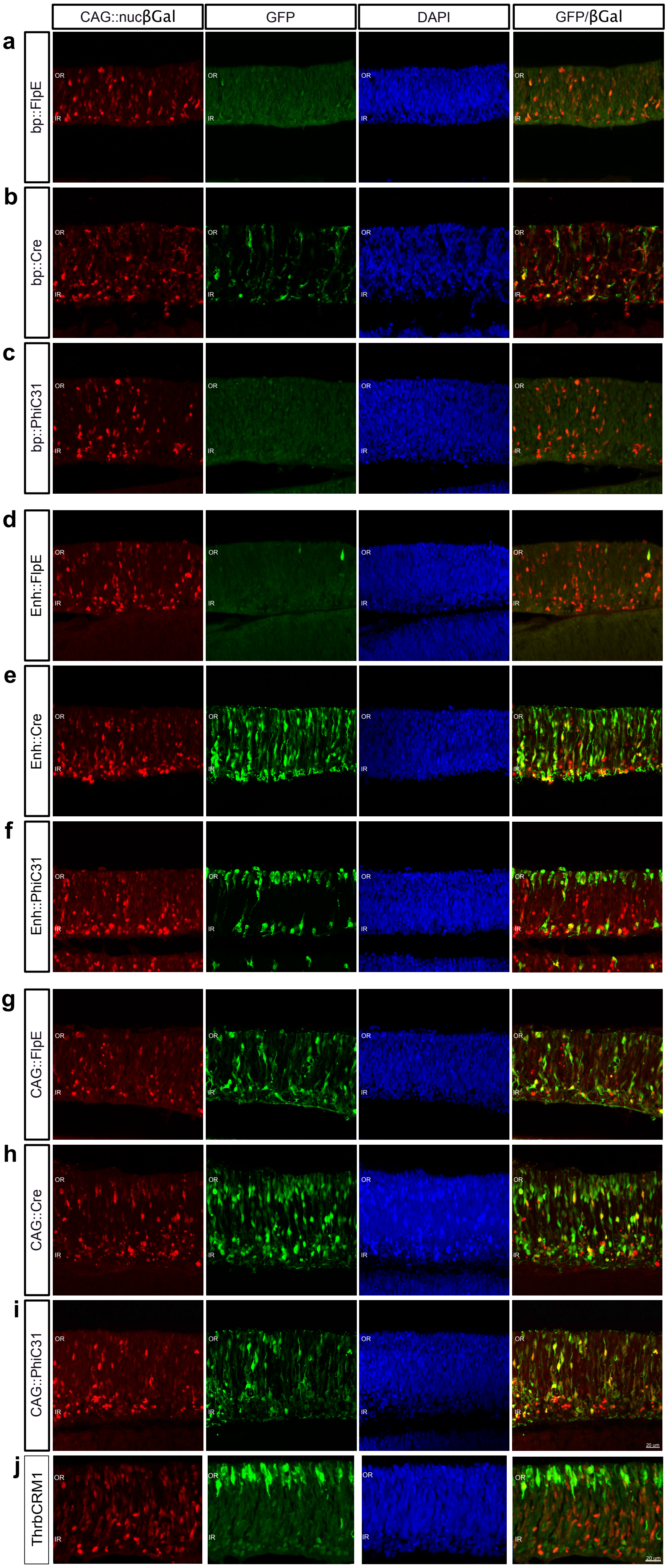

### Supplementary Figure S2

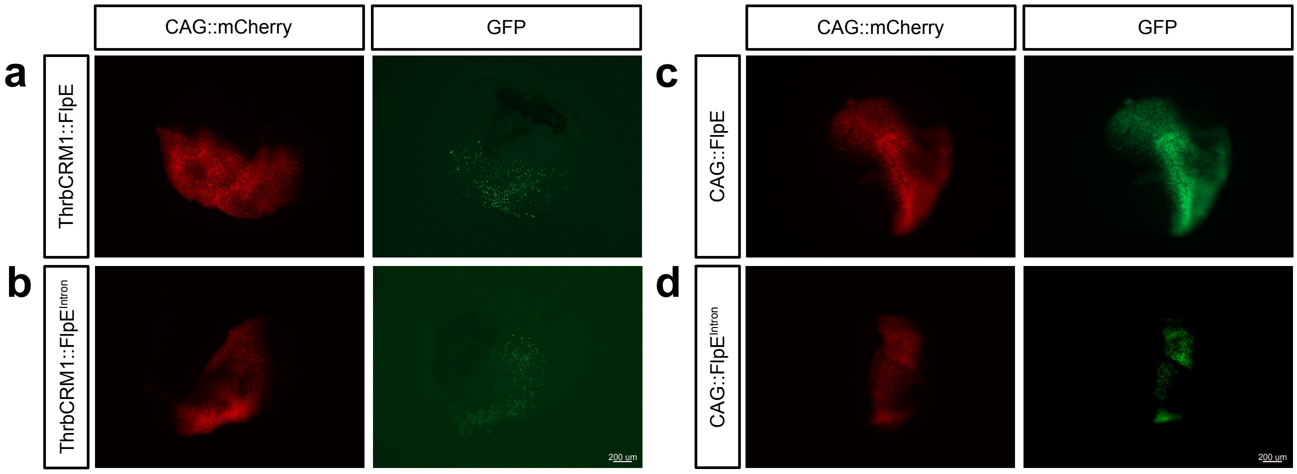


### Supplementary Figure S3

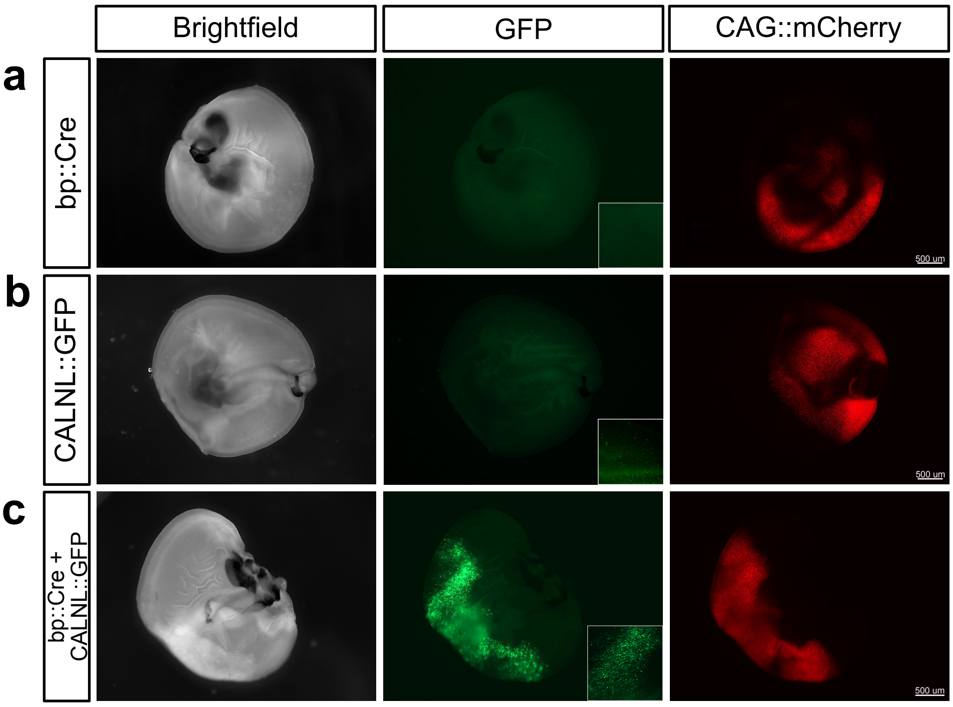

### Supplementary Figure S4

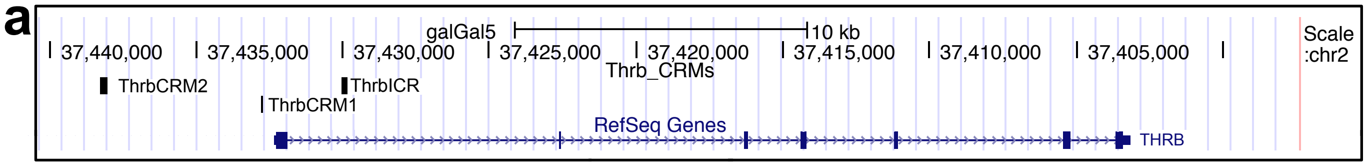


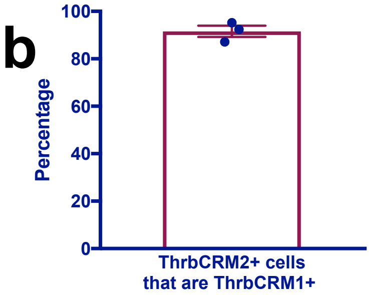


### Supplementary Figure S5

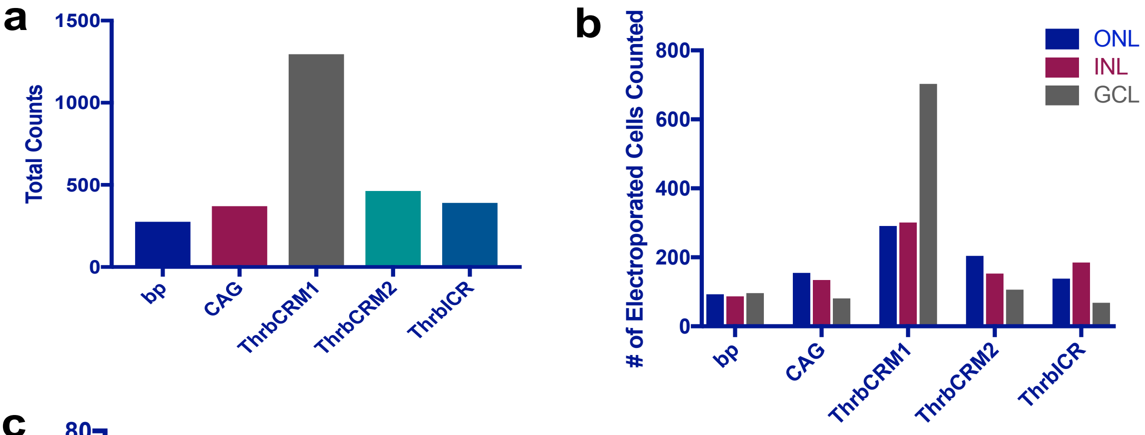


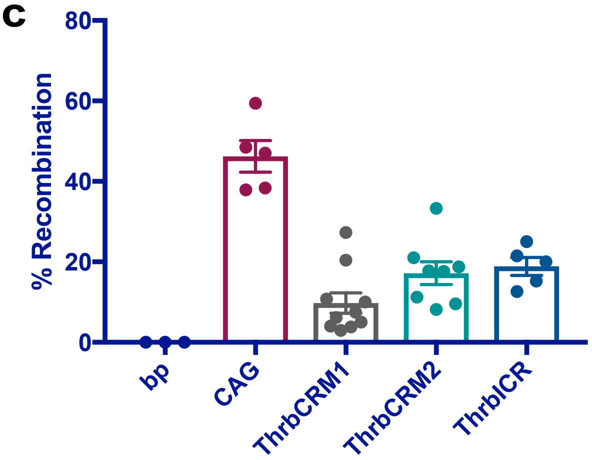

### Supplementary Figure S6

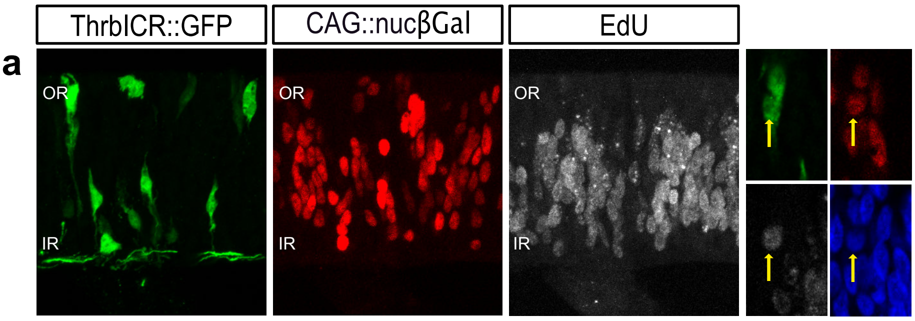


### Supplementary Figure S7

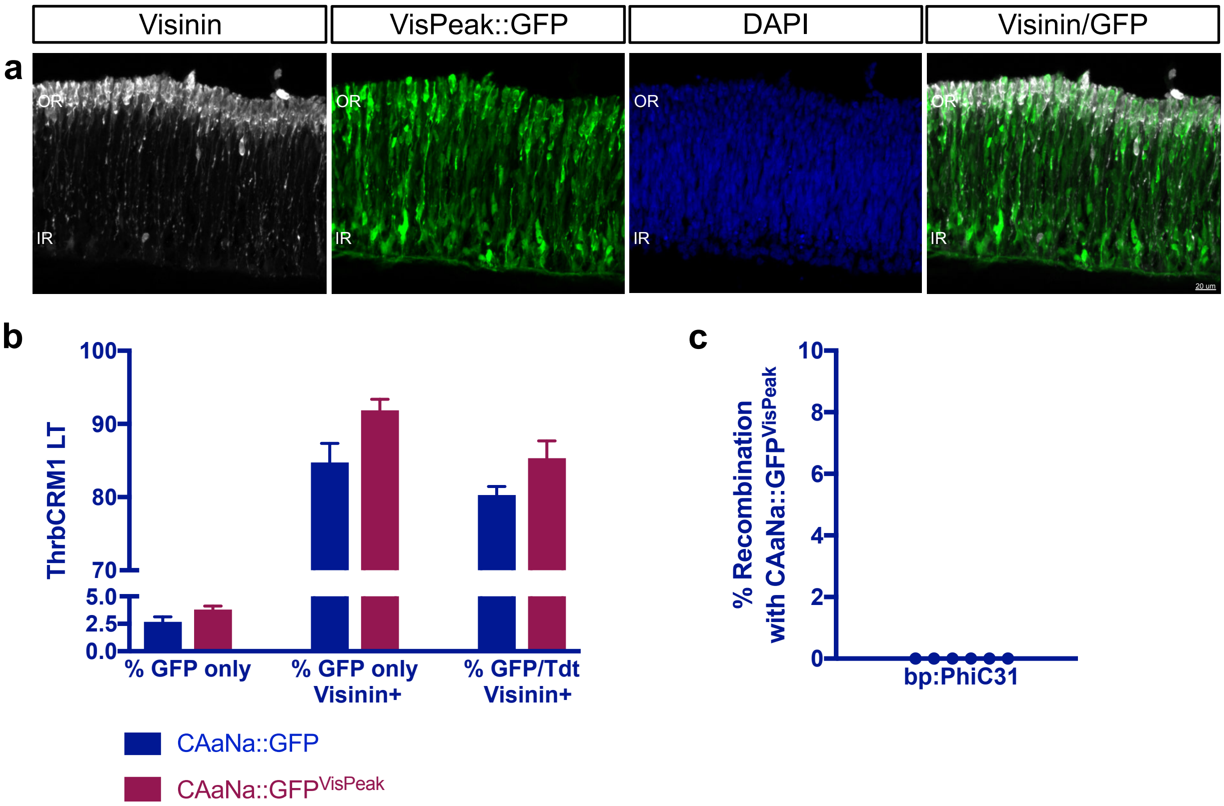

Supplementary Table S1. Primer Sequences

| | |
|---|---|
| **Bolded nucleotides correspond to restriction sites, italicized nucleotides correspond to attB or attP sequences | |
| PhiC31 Xma1-tagged forward primer | 5’ AG**CCCGGG**ACCATGGATACCTACGCCGGAGCC 3’ |
| PhiC31 BsrG1-tagged reverse primer | 5’ AG**TGTACA**TCACACTTTCCGCTTTTTCTT 3’ |
| attPlongNeoF1 | 5’ *TTTGAGTTCTCTCAGTTGGGGGCGTAG*TCGGATTTGATCTGATCAAGAG 3’ |
| attBLongNeoR1 | 5’ *CCAAGGGCACGCCCTGGCACCCGCACCGCGGCTTCGAG*ACGCGTTCGGATTTGATCCAG 3’ |
| attPLongNeoF2 | 5’ GA**CTCGAG***GTGCCCCAACTGGGGTAACCTTTGAGTTCTCTCAGTTGGGGGCG* 3’ |
| attBLongNeoR2 | 5’ AG**CTCGAG***GATGGGTGAGGTGGAGTACGCGCCCGGGGAGCCCAAGGGCACGCCCTGGC* 3’ |
| attP total sequence | 5’ GTGCCCCAACTGGGGTAACCTTTGAGTTCTCTCAGTTGGGGGCGTAG 3’ |
| attB total sequence | 5’ CTCGAAGCCGCGGTGCGGGTGCCAGGGCGTGCCCTTGGGCTCCCCGGGCGCGTACTCCACCTCACCCATC 3’ |
| VisPeak Forward primer | 5’ TAGCGCTCCTTAATGACCGG 3’ |
| VisPeak Reverse primer | 5’ GCGGGATTAAAGCGGTCGT 3’ |
| VisPeak Bcu1-tagged forward primer | 5’ ATT**ACTAGT**TAGCACTCCTTAATGACCGG 3’ |
| VisPeak Reverse primer | 5’ TTC**ACTAGT**GATTGCGGGATT 3’ |

Supplementary Table S2. FlpE/ACTB intron Geneblock sequence

| |
|---|
| **Shaded nucleotides correspond to Swa1 and Age1 restriction sites, respectively. Underlined nucleotides refer to conserved exon sequences, bolded nucleotides correspond to the ActinB intron sequence. |
| CAGTTCGAATCATCGGAAGAAGCAGATAAGGGAAATAGCCACAGTAAAAAAATGCTTAAAGCACTTCTAAGTG<br>AGGGTGAAAGCATCTGGGAGATCACTGAGAAAATACTAAATTCGTTTGAGTATACCTCGAGATTTACAAAAACA<br>AAAACTTTATACCAATTCCTCTTCCTAGCTACTTTCATCAATTGTGGAAGATTCAGCGATATTAAGAACGTTGATC<br>CGAAATCATTTAAATTAGTCCAAAATAAGTATCTGGGAGTAATAATCCAGTGTTTAGTGACAGAGACAAAGACA<br>AGCGTTAGTAGGCACATATACTTCTTTAGCGCAAGGGGTAGGATCGATCCACTTGTATATTTGGATGAATTTTTG<br>AGGAACTCTGAACCAGTCCTAAAACGAGTAAATAGGACCGGCAATTCTTCAAGCAACAAACAGG**TAAGTGACC**<br>**TGTTACTTTGGGAGTGGCAAGCCTGGGGTTTTCTTGGGGATCGATGCCGGTGCTAAGAAGGCTGTTCCCTTCC**<br>**A**CAGGAATACCAATTATTAAAAGATAACTTAGTCAGATCGTACAACAAGGCTTTGAAGAAAAATGCGCCTTATC<br>CAATCTTTGCTATAAAGAATGGCCCAAAATCTCACATTGGAAGACATTTGATGACCTCATTTCTGTCAATGAAGG<br>GCCTAACGGAGTTGACTAATGTTGTGGGAAATTGGAGCGATAAGCGTGCTTCTGCCGTGGCCAGGACAACGTA<br>TACTCATCAGATAACAGCAATACCTGATCACTACTTCGCACTAGTTTCTCGGTACTATGCATATGATCCAATATCA<br>AAGGAAATGATAGCATTGAAGGATGAGACTAATCCAATTGAGGAGTGGCAGCATATAGAACAGCTAAAGGGT<br>AGTGCTGAAGGAAGCATACGATACCCCGCATGGAATGGGATAATATCACAGGAGGTACTAGACTACCTTTCAT<br>CCTACATAAATAGACGCATAGGACCGGTGGAACAAAAAC |
